# Anticipating change: the impact of simulated seasonal heterogeneity on heat tolerances along a latitudinal cline

**DOI:** 10.1101/2023.02.19.529162

**Authors:** Jared Lush, Carla M. Sgrò, Matthew D. Hall

## Abstract

An understanding of thermal limits and variation across geographic regions is central to predicting how any population may respond to scenarios of global change. Latitudinal clines, in particular, have been used in demonstrating that populations can be locally adapted to their own thermal environment and, as a result, not all populations will be equally impacted by an increase in temperature. But how robust are these signals of thermal adaptation to the other ecological challenges that animals commonly face in the wild? Seasonal changes in population density, food availability, or photoperiod are common ecological challenges that could disrupt patterns of thermal tolerance along a cline if each population differentially used these signals to anticipate future temperatures and adjust their thermal tolerances accordingly. In this study, we aimed to test the robustness of a cline in thermal tolerance to simulated signals of seasonal heterogeneity. Experimental animals were derived from clones of the Australian water flea, *Daphnia carinata*, sampled from nine populations along a latitudinal transect in eastern Australia. We then factorially combined summer (18h light, 6h dark) and winter (6h light, 18h dark) photoperiods with high (5 million algal cells individual^-1^ day^-1^) and low (1 million algal cells individual^-1^ day^-1^) food availabilities, before performing static heat shock assays and recording knockdown times as a measure of thermal tolerance. In general, higher food availably led to an increase in thermal tolerances, with the magnitude of increase varying by clone. In contrast, summer photoperiods led to rank order changes in thermal tolerances, with heat resistance increasing for some clones, and other decreasing for others. Heat resistance, however, still declined along the latitudinal cline, irrespective of the manipulation of seasonal signals, with northern clones always showing greater thermal resistance, and that this was most likely driven by adaptation to winter thermal conditions. While photoperiod and food availability can clearly shape thermal tolerances for specific clones or populations, they are unlikely to be used to anticipate future temperatures, and thus observed clines in heat resistance will remained robust to these forms of seasonal heterogeneity.

## Introduction

Global change has led to the gradual increase in both mean temperatures and extreme climatic events and is forecast to continue into the future (Schär et al. 2004; Rahmouf and Coumou 2011; Buckley and Huey 2016b). An understanding of thermal limits and variation amongst geographic regions is central to predicting how any population may respond to scenarios of global change (Hoffmann et al. 2002; Sgrò et al. 2010; Hoffmann and Sgró 2011). Latitudinal clines, in particular, have been used in demonstrating that populations are commonly locally adapted to their own thermal environment and, as a result, not all populations will be equally impacted by an increase in temperature (Kellermann et al. 2006; Sgrò et al. 2010; Yampolsky et al. 2013). Accurate predictions for a population’s response to global change, however, depend on the robustness of thermal limit clines to other sources of environmental variation that regularly act to shape the life-history and physiology of individuals at each location (Brinkhof and Cave 1997; Lepage et al. 1998; Moran et al. 2016).

Within any population, individuals will be exposed to a range of environmental conditions throughout the year that can modify their tolerance to heat stress, including stochastic variation in food supply (Nguyen et al. 2017), sudden heat waves (Buckley and Huey 2016a) or the presence of competitors or parasites (Hector et al. 2019; Thawley et al. 2019). These biotic and abiotic factors can disrupt the predictable change in thermal limits that is the hallmark of latitudinal clines, by either shifting the average thermal tolerance of all populations equally, or by inducing population-specific responses that can reduce or even completely erode the previously established clines (Hector et al. 2019; Thawley et al. 2019). Exposure to other stressors, such as invasive species (Thawley et al. 2019) or pathogens (Hector et al. 2019), for example, have been shown to negate the increase in upper thermal limits that is expected as a population moves closer to the equator. In contrast, direct exposure to both gradual and abrupt increases in temperature (Jørgensen et al. 2019) are well-established mechanisms by which individuals can acclimate and harden to exposed thermal stress and improve thermal resistance for the future (Hoffmann and Parsons 1994; van Heerwaarden et al. 2016).

Seasonal changes in population density, food availability, or photoperiod (Bradshaw and Holzapfel 2007; Jiang et al. 2014) also offer the opportunity to anticipate future temperatures and adjust thermal tolerances accordingly (Manenti et al. 2021). Many bird species, for example, use photoperiod for timing long migrations or breeding periods with peak resource (Dawson et al. 2001; Cohen et al. 2020). Whether or not, however, species also have the capacity to alter their thermal resistance in response to signals associated with the changing of seasons remains largely unknown. Recent studies have attempted to determine the impact of seasonal variation on knockdown resistance by manipulating photoperiod as a proxy for season. Few studies have found an effect of photoperiod alone, but this may be due to the method of applying treatments, with one study showing that daily increasing or decreasing photoperiods effect different stress resistances, albeit species dependent (Manenti et al. 2021). We do, however, have evidence for male-specific reductions in knockdown times during summer light conditions (Bauerfeind et al. 2014) and population dependent increases in CT_max_ due to increasing photoperiod (Moghadam et al. 2019). While these studies highlight the potential for signals of seasonality, such as photoperiod, to modulate a population’s thermal tolerance, and to do so in a population or genotype specific manner, they do not necessarily capture all aspects of seasonal change or whether all populations along a cline will respond in a similar manner.

In this study, we aim to identify a latitudinal cline for thermal tolerance in the Australian water flea, *Daphnia carinata* and then determine how robust any cline may be to signals of seasonal heterogeneity. *Daphnia* are ideal for testing both the robustness of clinal variation in thermal limits to seasonal change, and the capacity of individuals to anticipate future conditions using the same signals. A mix of photoperiod, food availability, and population density are commonly used as signals for determining life-history and behaviour (Roulin et al. 2013; Jiang et al. 2014; Gust et al. 2019; Lever et al. 2021). For Australian *Daphnia*, for example, longer photoperiods are congruent with harsh summer conditions that lead to seasonal population crashes, a signal which is used to increase the number of sexually produced eggs, relative to asexual reproduction (Lever et al. 2021). *Daphnia* are also sensitive to changes in the thermal environments, with temperature driving key ecological processes such as migration (Burns 2013), growth (Martínez-Jerónimo 2012) and fecundity (Wojtal-Frankiewicz 2012). Indeed, temperature is a strong driver of evolution, with not only evidence of adaptation to population’s local thermal environment (Williams et al. 2012; Yampolsky et al. 2013) but also that this adaptation is sensitive to other factors such as competition (De Meester et al. 2011). However, while *Daphnia* have yet to be used in a study combining exploring links between non-thermal seasonal cues and thermal tolerance.

Here, we integrated a manipulation of environmental cues associated with seasonality into the study of thermal limits along a latitudinal cline. We expected that a latitudinal cline for thermal tolerance would be evident, with populations closer to the equator exhibiting larger resistance to knockdown, as they would be locally adapted to higher temperatures (Hoffmann et al. 2002). We aimed to test if different clones can use seasonal change to improve future thermal tolerance, by manipulating two key variables that vary significantly with season in our system – photoperiod and food availability (Lever et al. 2021). We expected that if thermal resistance of clones was sensitive to seasonal signals, that clones from latitudes with large variation in seasonal photoperiod would be more likely to use these seasonal cues (Bradshaw and Holzapfel 2007), with food availability amplifying any effect due to the condition-dependence of thermal tolerances. To complement the above, we also used microclimate data available for each population to explore the environmental factors potentially underlying adaptation to thermal tolerance along a cline in *D. carinata*. We expected that winter conditions may drive any local adaptation, as our populations are largely present during winter, as evident in previous research on European *Daphnia* (Seefeldt and Ebert 2019).

## Methods

### Study system

Experimental animals were derived from nine clones (PLD, BWD, BLD3, WWD2, YYD, OZD, DC28, CMD and LBD, see Figure 1 and Table S1) of *Daphnia carinata* sampled from populations varying in latitude along a cline from Tamworth to Geelong in south-eastern Australia (Drapes et al. 2021; Lever et al. 2021). Clones were maintained under standard laboratory conditions, stored at 20 °C and under a 16:8 L:D light regime. From the stock animals, individual females presenting eggs were first isolated in 70-ml glass jars containing 40 mL of Artificial Daphnia Medium (ADaM, Klüttgen et al. 1994, modified after Ebert et al. 1998). Daphnia were maintained individually for three generations to reduce maternal effects, before isolating experimental animals from clutches three to seven, correlating with block numbers one to four.

**Figure 1:**
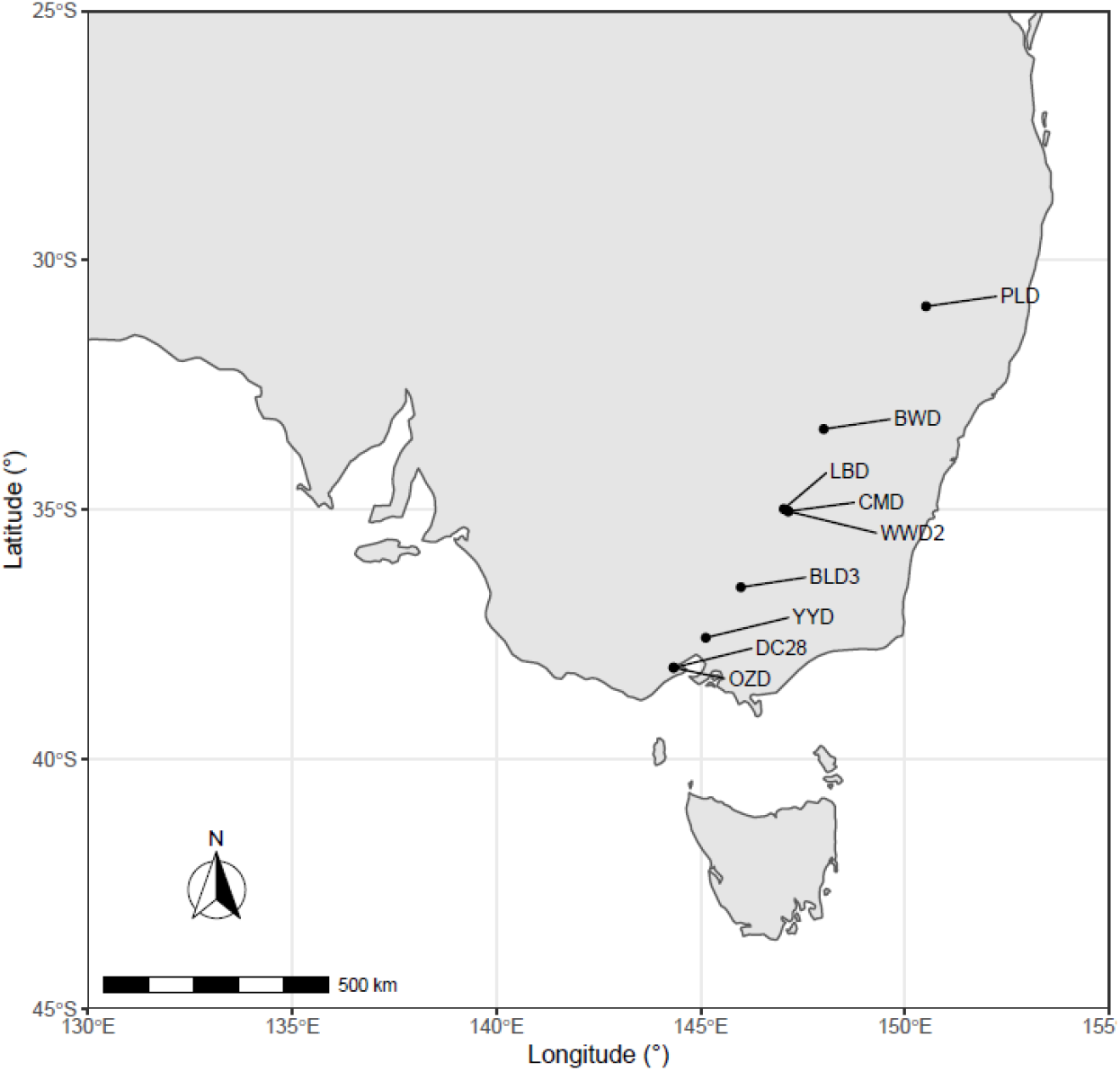
Map of South-eastern Australia showing locations sourced for *Daphnia carinata*

**Figure 2:**
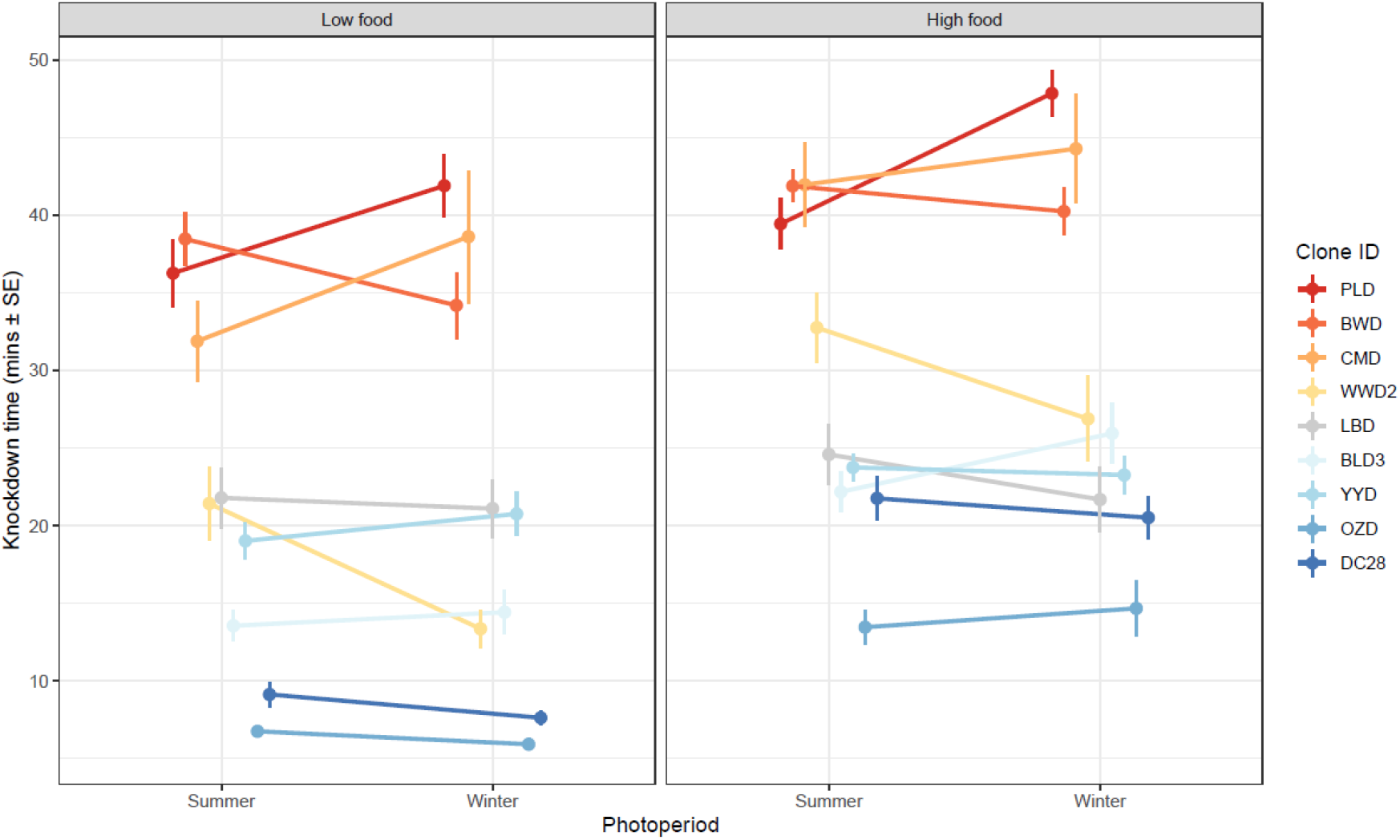
Reaction norm for the effect of photoperiod and food availability on knockdown time for each host genotype of *Daphnia carinata*. High food treatment consisted of 5 million green algal cells daily, low food was 1 million green algal cells daily. Summer photoperiods consisted of an 18:6-hour photoperiod, while winter photoperiods consisted of a 6:18 hour photoperiod. Points displayed are mean values (± SE) for knockdown times at 37 °C. Clone IDs are displayed in the legend and are ordered by latitude, with populations from lower latitudes (°S) at the top (red) and higher latitude populations at the bottom (blue).

### populations

#### Basic experimental design

Seasonal treatments were created by manipulating photoperiods and food availability. In addition to increasing photoperiods leading into summer, natural populations see an increase in food availability due to increases in abundance of algal species (Moss et al. 2003), but also a decrease in competition due to lower population densities (Barry 1997). To mimic this situation, we factorially combined summer (18h light, 6h dark) and winter (6h light, 18h dark) photoperiods (as in: Lever et al. 2021), with high (5 million algal cells individual^-1^ day^-1^) and low (1 million algal cells individual^-1^ day^-1^) food availabilities (as in: Lever et al. 2021), resulting in four total treatment groups. Each treatment contained 40 experimental animals (9 genotypes x 2 food qualities x 2 seasons = 40 × 40 individuals = 1440 total animals). Animals were maintained under standard conditions in controlled climate chamber (20°C, medium changed twice weekly) and were fed daily with green algae (*Scenedesmus sp*.) until static heat shock assays were performed. Within the climate chambers, the location of animals was shuffled daily to minimise any positional effects.

#### Static heat shock assays

Static heat shock assays were performed to determine the time to immobilisation (*T*_*imm*_) as a measure of thermal tolerance, herein referred to as knockdown time (as in: Yampolsky et al. 2014). Animals were assayed at 20 ± 1 days old. Assays were performed in 70 L glass aquariums, using a thermoregulator to maintain temperatures at 37 °C with constant agitation and bubblers to maintain oxygen saturation throughout the assay. Animals were kept inside 5 ml glass vials, covered by a mesh lid to prevent oxygen depletion within the vial (as per Hector et al. 2019). Knockdown times were scored beginning from when the vials enter the assay until individuals cease all visible movement, including filtering (Yampolsky et al. 2014; Hector et al. 2019). Individuals from each population were randomly selected for each assay. Each assay could contain 48 animals, and as such 4 – 5 animals from each population were present in each assay. Assays for each block were performed over two days, with four assays per day. Immediately following the static heat shock assays, the body size of each individual was measured via stereo microscope from the individuals eye to the base of the tail (see Laidlaw et al. 2020).

#### Statistical analyses

All analyses were performed using R (v 4.0.5, 2021; www.R-project.com). We first assessed the impact of seasonal change on each population’s thermal response via a linear mixed-effect model as implemented via the lme4 package (Bates et al. 2015). This model included knockdown time as the response variable, and photoperiod (summer and winter), food availability (high and low) and *Daphnia* genotype (9 levels), as interacting fixed effects. Body size was included as a covariate while experimental block was used as a random effect. We then related how changes in thermal tolerance may relate to the location of each *Daphnia* clone along the latitudinal cline. To do so we used a two-factor analysis of covariance with photoperiod and food availability as fixed effect factors and latitude (°S) as the interacting covariate, as implemented as a linear mixed-effect model with experimental block as a random effect again.

To complement the above analyses, we used microclimate data available for each population to explore the environmental factors potentially underlying adaptation to thermal tolerance along a cline in *D. carinata* (Bramer et al. 2018). Microclimate data was obtained for 10 km^2^ areas at the source GPS locations for each populations (Databases originally sourced from New et al. 2002; data accessed through http://niche-mapper.com/apps/climate), including average monthly min and max temperatures, precipitation data, wind speed, altitude, cloud cover and humidity (see Table S1). Data was summarised to obtain average values of each variable for winter and summer months, as populations do not persist year-round. Independent variables were then mean scaled and centred before analysis.

To fit the environmental data to knockdown times, we used the dredge function from the ‘MuMin’ package (Barton 2022). The package initially created linear models with knockdown times as a response and containing all combinations of up to 5 explanatory variables from the19-total available (Table S2), including the experimental treatments (food availability by photoperiod) as a required fixed effect. Model selection compared all model permutations by their Akaike information criterion (AIC). Models within AIC < 2 from the best fitting model would be averaged to obtain effect estimates of each parameter.

## Results

We first considered how seasonal variation affected upper thermal limits by manipulating food availability and photoperiod and then performing static heat shock assays. We found that knockdown times varied independently with both food availability and photoperiod, but that any effect depended on the *Daphnia* clone (i.e., two-way interaction terms, Table 1a). The nature of the clone-specific responses, however, differed between the two environmental manipulations. Reducing the availability of food, for example, led to a 6-minute, on average, decrease in knockdown times, with clones differing in the magnitude of effect, for example for DC28, knockdown times decreased by ∼12 minutes with diet, while LBD decreased by less than 3 minutes (Figure 1). In contrast, in response to changes in photoperiod the *Daphnia* clones varied in both the direction and magnitude of change in their knockdown times, leading to rank-order changes in the performance of clones in any period. This is evident by the increase in knockdown times with photoperiod shown by BWD, as contrasted to the decrease that PLD exhibited, causing a change in the order of best performing populations depending on the length of photoperiod.

**Table 1:** Linear mixed-effect models for the effects of a) size, photoperiod (winter light length and summer light length), food availability (high food, low food), and host genotype on knockdown times during a 37 °C static heat shock assay. b) size, photoperiod (winter light length and summer light length), food availability (high food, low food) and original latitude of host population on knockdown times during a 37 °C static heat shock assay. Block was a random effect, for a) σ^2^= 5.55, σ = 2.36, for b) σ^2^ = 6.17, σ = 2.45.

| Term | df | $\chi^2$ | p-value |
| --- | --- | --- | --- |
| *(a) Impact of host population* |  |  |  |
| Food availability | 1 | 7.23 | 0.007** |
| Photoperiod | 1 | 0.01 | 0.154 |
| *Daphnia* clone | 8 | 269.46 | < 0.001*** |
| Photoperiod x *Daphnia* clone | 8 | 22.55 | 0.004** |
| Food availability x *Daphnia* clone | 8 | 17.28 | 0.027* |
| Photoperiod x Food availability x *Daphnia* clone | 8 | 4.75 | 0.784 |
| Photoperiod x Food availability | 1 | 0.96 | 0.664 |
| Size | 1 | 1.33 | 0.249 |
| *(b) Impact of population location* |  |  |  |
| Food availability | 1 | 2.11 | 0.146 |
| Photoperiod | 1 | 2.73 | 0.098# |
| Latitude | 1 | 148.53 | < 0.001*** |
| Photoperiod x Latitude | 1 | 2.64 | 0.104 |
| Food availability x Latitude | 1 | 4.07 | 0.044* |
| Food availability x Photoperiod | 1 | 0.52 | 0.471 |
| Food availability x Photoperiod x Latitude | 1 | 0.46 | 0.499 |
| Size | 1 | 3.77 | 0.052# |

To test if the variation in knockdown times could be related to the process of local adaptation occurring along a latitudinal cline, we analysed changes in knockdown time in response to treatment effects and source latitudes of the genotypes. We found a negative relationship between latitude (°S) and knockdown times (Table 1b; Figure 3). Clones derived from populations closer to the equator tended to have higher knockdown times than populations further away, confirming a latitudinal cline in thermal resistance for the latitudes studied. The relationship between latitude and knockdown times was marginally affected by the manipulation of seasonal signals, as captured by the significant food availability by latitude interaction (Table 1b). Reductions in food availability caused a 16 percent increase in the rate at which knockdown times decline with distance from the equator (slope change of 3.78 ± SE min per degree south to 4.40 ± SE per degree south).

**Figure 3.**
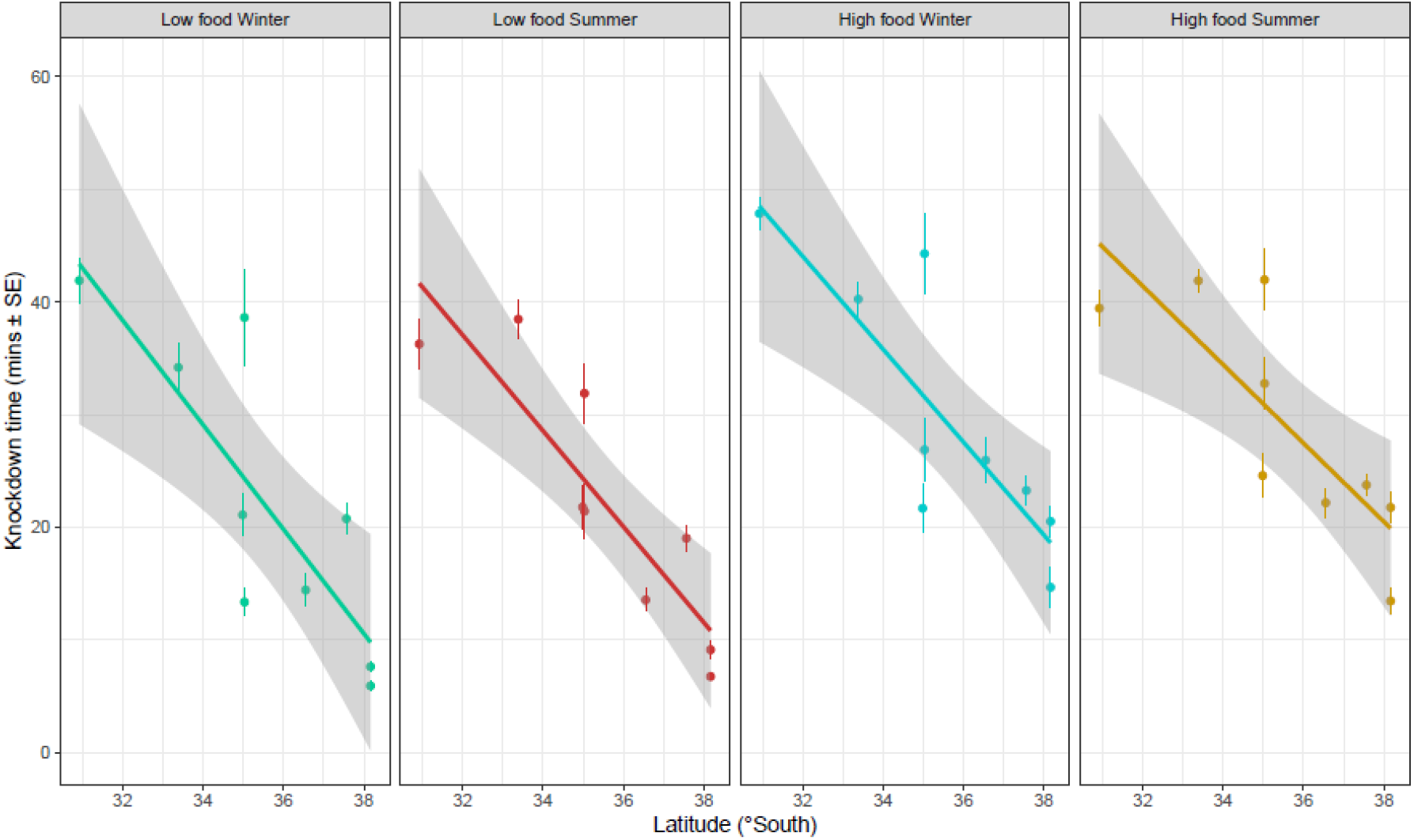
Knockdown times at 37 °C for populations along a latitudinal cline (°S) for each food availability and photoperiod treatment of *Daphnia carinata*. High food availability treatment consisted of five million green algal cells daily, low food availability one million green algal cells daily. Summer photoperiod treatments consisted of an 18:6-hour photoperiod, whilst winter photoperiod was 6:18 hour. Values shown are mean (± SE) knockdown times for each clone and the predicted trend across latitude (± CIs).

To understand the potential environmental drivers of the cline in knockdown times, we obtained fine-scale microclimate data and modelled predictors against knockdown times. The best fitting model contained the variables wind speed, average high temperature during winter months (AHT winter) and average monthly min temperature (ALT), as well as the fixed treatment effects (Figure 4). Both predictors pertaining to temperature were positively correlated with thermal tolerance, with populations that experienced higher winter AHT and higher ALT associated having elevated knockdown times (Figure 4). Conversely, the average wind speed of each location was negatively associated with knockdown times (Figure 4). Importantly, only the temperature predictors that were conducive of winter were present in the significant model, not changes in photoperiod. This suggests that seasonal temperatures in combination with wind speeds may be mostly responsible for local adaptation of thermal resistance, with warmer temperatures and lower wind speeds during the winter month potentially leading to the evolution of greater thermal tolerance.

**Figure 4.**
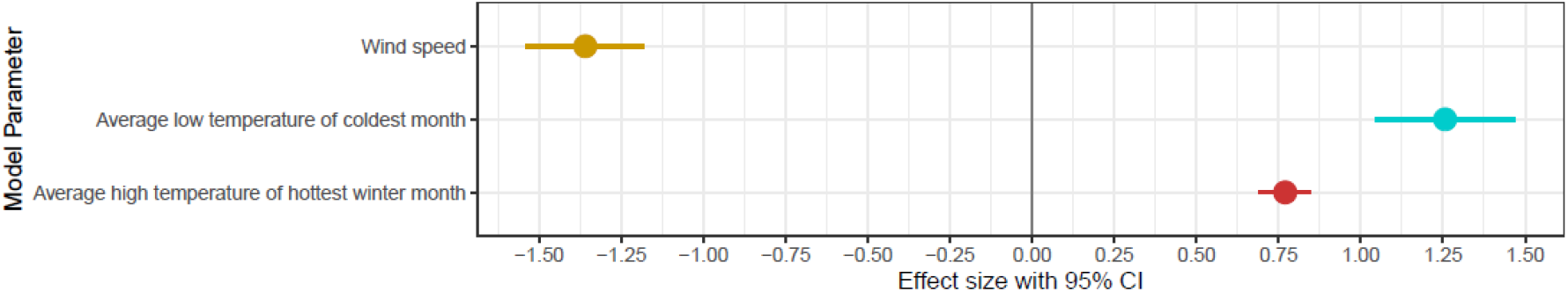
Effects of environmental variables on knockdown times of *Daphnia carinata*. Effect size estimates are scaled and centred and from the best fitting model (lowest AIC value). Error bars indicate 95% confidence intervals. Parameters missing from the figure were among variables that were omitted from the best fitting model, and thus do not contain estimates. Photoperiod and food availability treatments were included in all models but omitted from figure.

## Discussion

The capacity of individuals and populations to anticipate future environmental condition and adjust their thermal tolerances accordingly is important for population persistence but may misconstrue our interpretation of clinal relationships. Here we analysed the thermal resistance of *D. carinata* sampled from along a latitudinal cline in response to two common seasonal cues. Initially, we proposed that populations may be able to use these signals of seasonal change, herein the increase in food and increase in photoperiod that signal the coming of summer, to prepare for future thermal conditions. Our results showed that both food availability and photoperiod indeed appear to influence the thermal tolerance of an individual, but their effects, not surprisingly, depended entirely on the clone of *Daphnia* used.

For photoperiod, we observed rank order changes in the reaction norms, with some clones performing better under summer light conditions and others performing better at winter light conditions. In contrast, low food availability led to a relatively consistent decrease in knockdown times across all clones, with only the magnitude of the effect differing, which is similar to what has been observed in other species where we see prolonged dietary restriction reducing thermal resistance with length of starvation (Nguyen et al. 2017; Manenti et al. 2018; Mulaudzi et al. 2022). The presence of these clone-specific responses rules out the possibility that all clones are uniformly using food availability and photoperiod signals to modify their thermal tolerances, and highlights how plastic thermal limits can be in response to the repeatable change that occurs each year, not just temperature changes that have been the focus of acclimation studies (Alford et al. 2012; Leclair et al. 2020).

To explore how the geographic location of each clone may shape its thermal tolerance in light of seasonal change, we modelled knockdown times against the latitude of sampled clone populations. We found clear evidence of a latitudinal cline in thermal knockdown times, with populations closer to the equator exhibiting greater thermal resistance. The clinal relationship was similar in direction and magnitude to what we know for general ectotherm species (Hoffmann et al. 2002; Sunday et al. 2011) and other *Daphnia spp* (Yampolsky et al. 2013; Seefeldt and Ebert 2019). For example, in *D. melanogaster* knockdown times decline at a rate of 0.21 minutes per degree south for heat shock assays at 39 °C (Hoffmann et al. 2002) or 0.15 minutes per degree south at 37 °C (Sgrò et al. 2010). Here, we observed a decrease of 3.78 minutes per degree south in high diet treatments or 4.40 minutes per degree south in low diet treatments, which corresponds with the relatively greater latitudinal variation in thermal resistance that is known to be a property of aquatic species when compared to terrestrial species (Sunday et al. 2011).

Driving these results is likely local adaptation to the thermal conditions experienced by each cline in winter (Figure 4). Across a wide collection of fine-scale microclimate data, both high and low winter temperatures remained some of the best predictors of variation in knockdown times amongst the clones. This is consistent with what we know about *D. carinata*, with populations expanding during cooler autumn, winter and spring conditions before declining during the extreme heat of summer (Vijayaraghavan 1970; Lund and Davis 2000; Burns 2013). By comparison, 22 European clones of *Daphnia* (*D. magna)* increase thermal resistance depending on their local thermal environments and are predicted to be locally adapted to the average high temperature during the hottest month of the year. (Yampolsky et al. 2013), although a similar trend to our study has been observed for a subset of ‘winter-active’ *D. magna* populations (Seefeldt and Ebert 2019). Given the limited ability to behaviourally mitigate warmer temperatures and limited gene-flow between patches (Hebert and Wilson 1994), it is not surprising that winter temperatures might be the main selective pressure behind thermal resistance in this system.

Despite observing strong changes to knockdown times due to changes in photoperiod and food availability at the clonal level, the relative changes in thermal tolerance along the latitudinal transect remained relatively unaffected (Figure 3). We initially hypothesised clones sourced further from the equator would be more likely to plastically adjust their phenotype in response to seasonal variation (Heideman et al. 1999; Śniegula et al. 2012). This, however, was not the case as the most significant photoperiod induced changes did not occur with clones and populations from the ends of the cline. Instead, a subtle increase in the strength of the clinal signal arose when animals were food limited, consistent with idea that thermal limits are condition dependent (Nyamukondiwa and Terblanche 2009; Bujan and Kaspari 2017). Nonetheless, the general robustness of the observed clinal relationship to these signals of seasonal change reinforces the utility of using clinal data for understanding and predicting the response of populations to climate scenarios (Deutsch et al. 2008; Kingsolver and Buckley 2017). As the difference in heat resistance between population outweighs any localised effects of seasonal change, they allow insight into the strength of local adaptation for thermal resistance and can assist in including within species heterogeneity in thermal tolerance based on latitude (Hoffmann and Sgró 2011; DeMarche et al. 2019). This is of particular importance as many models assume relatively constant thermal tolerance across time, which if our results had proved not to be true, would provide inaccurate outcomes that generalise a species response across its entire range (Kingsolver and Buckley 2017; DeMarche et al. 2019).

In conclusion, we observed clone-specific effects of photoperiod as well as general increases in knockdown times due to increases in food availability, signifying the important influence that seasonal change, beyond just temperature, can have on stress resistance. The strength of the effect attributed to local adaptation to winter temperatures, however, overshadowed any seasonal effects when including spatial data in the analyses. Our results thus contribute to a growing awareness for the ecological conditions under which latitudinal clines in heat resistance will remain robust, or not, adding to the studies which have shown how warming generally increases the heat tolerance of most populations equally (Hoffmann and Parsons 1994; van Heerwaarden et al. 2016), whereas immune activation (Hector et al. 2019) or invasive species (Thawley et al. 2019) instead erodes any clinal signal. This is an important step going forward, providing more confidence in using clinal data of thermal resistance in predicting how locally adapted populations will respond to future climate scenarios.

## Acknowledgements

This work was supported by funding from Monash University and the Australian Research Council.

