## Supplemental Tables for "Anticipating change: the impact of simulated seasonal heterogeneity on heat tolerances along a latitudinal cline"

**Table S1.** Geographic location information for each *Daphnia carinata* clonal line.

| Population | Location | Latitude | Longitude | Altitude |
|---|---|---|---|---|
| **PLD** | Tamworth (NSW) | -30.9222 | 150.5181 | 296 |
| **BWD** | Lake Forbes (NSW) | -33.3831 | 148.0119 | 240 |
| **LBD** | Wagga Wagga (NSW) | -34.9833 | 147.0361 | 178 |
| **CMD** | Wagga Wagga (NSW) | -35.0297 | 147.1439 | 173 |
| **WWD2** | Wagga Wagga (NSW) | | | 173 |
| **BLD3** | Benalla (VIC) | -36.553401 | 145.984488 | 173 |
| **YYD** | Yan Yean (VIC) | -35.0297 | 147.1439 | 178 |
| **DC28** | Geelong (VIC) | -38.165702 | 144.338144 | 12 |
| **OZD** | Geelong (VIC) | -38.165702 | 144.338144 | 12 |

**Table S2.** Environmental variables used for the microclimate model.

| Variable | Sourced from |
|---|---|
| Summer day length | Geosphere package, v1.5-18 2022 |
| Winter day length | Geosphere package, v1.5-18 2022 |
| Yearly photo period variation | |
| Average high temperature of hottest month (AHT) | Niche Mapper, http://niche-mapper.com/apps/climate |
| AHT (summer) | |
| AHT (winter) | |
| Average low temperature of coldest month (ALT) | Niche Mapper, http://niche-mapper.com/apps/climate |
| ALT (summer) | |
| Average high rainfall (AHR) | Niche Mapper, http://niche-mapper.com/apps/climate |
| AHR (summer) | |
| AHR (winter) | |
| Average low rainfall (ALR) | Niche Mapper, http://niche-mapper.com/apps/climate |
| ALR (summer) | |
| ALR (winter) | |
| Wind speed | Niche Mapper, http://niche-mapper.com/apps/climate |
| Cloud cover | Niche Mapper, http://niche-mapper.com/apps/climate |
| Altitude | Elevatr package, v0.4.2 2022 |
| Treatment group | |
| Body size | Experimentally measured |
